# Dopamine serves as an independent gain regulator to shape striatal control of movement

**DOI:** 10.64898/2026.08.16.745105

**Authors:** Jingheng Zhou, Riley M. Harper, Amy B. Papaneri, Guohong Cui, Julieta E. Lischinsky

## Abstract

Dopamine (DA) drives locomotion by modulating striatal activity in the dorsolateral striatum (DLS). How DA regulates motor function on subsecond timescales remains poorly understood. To address this question, we performed spectrally resolved triple-color fiber photometry, allowing us to simultaneously monitor DA, glutamatergic inputs, and pathway-specific spiny projection neuron (SPN) activity in freely moving mice during spontaneous behaviors as measured by unsupervised behavior quantifications. We showed that DA dynamics were temporally distinct from spontaneous locomotor kinematics or behavioral states. Instead, peri-event DA levels predicted SPN input-output (I/O) efficacy, defined as SPN activity relative to glutamatergic input, with opposite relationships across pathways: higher DA predicted enhanced efficacy in direct-pathway SPNs but reduced efficacy in indirect-pathway SPNs. Paired with unsupervised behavior quantification, elevating extracellular DA with methylphenidate, a DA re-uptake inhibitor, shifted direct-pathway SPNs toward higher I/O gain and indirect-pathway SPNs toward lower neuronal activity outputs, while biasing behavior toward selected mobile and turning states. Reserpine administration, which mediates DA vesicular depletion, resulted in the opposite efficacy shifts and increased occupancy of immobile states. Together, these findings support that DA does not simply encode spontaneous movement, but acts as a pathway-specific gain controller that dynamically tunes the transformation of glutamatergic input into SPN output to bias locomotor-state transitions *in vivo*.

## Main

Neurotransmitter release at synapses is essential for driving postsynaptic responses and enabling signal transmission throughout the nervous system. In contrast to classical neurotransmitters, neuromodulators adjust neuronal and synaptic properties, thereby reshaping how circuits process information^1^. Among neuromodulators, dopamine (DA) in the striatum is the most extensively studied because of its role in motor control and learning, as well as reward and motivation^2–5^. Striatal spiny projection neurons (SPNs) integrate excitatory glutamatergic inputs from cortex and thalamus with dense dopaminergic innervation from the midbrain nigrostriatal pathway^6^. SPNs comprise two roughly equal populations: direct-pathway SPNs (dSPNs), which project to basal ganglia output nuclei, and indirect-pathway SPNs (iSPNs), which project to intermediate nuclei, collectively forming the canonical direct and indirect pathways of the basal ganglia^2,7^. Evidence from brain slices supported that DA increases dSPN excitability by enhancing cAMP production and protein kinase A (PKA) activity via G_s_-coupled D1 DA receptors (D1R), whereas DA decreases iSPN excitability by reducing cAMP and suppressing PKA through G_i/o_-coupled D2 DA receptors (D2R)^2–4,8–13^. Although the mechanisms of dopaminergic neuromodulation have been well studied in acute brain slices, concerns remain recently on how faithfully these processes operate in freely behaving animals. *In vivo* studies observed that 1) transiently altered DA levels by optogenetic active or inhibit DA neurons, led to only weak changes in striatal activity^14^, 2) no impairments in basic motor functions were detected in mice where phasic DA release was disrupted^15^, indicating that DA dynamics might play a minor role in the subsecond modulation of striatal dynamics, and are dispensable for movement. Therefore, the precise role of DA modulation on striatal activity for motor control remains elusive. To address this, in the present study we leveraged Cre transgenic mice lines^16,17^, *in vivo* imaging methods^18–20^ and genetically encoded fluorescent indicators^21–23^, and performed triple-color recordings to measure simultaneous DA and glutamate signaling together with SPN activity in freely moving mice. By integrating unsupervised behavior classification with pharmacological manipulations of DA, we investigated how DA shifts input-output efficacy of SPNs to reshape locomotor behavior.

## Results

### Striatal dopamine lacks movement tuning

To determine how striatal DA, glutamate, and SPN activity are coordinated during spontaneous locomotion, we co-expressed the DA indicator GRAB–gDA3m (gDA3m)^21^, the glutamate indicator SF–Venus–iGluSnFR.A184S (Venus–iGluSnFR)^22^ and the calcium indicator 2.0–sRGECO (sRGECO)^23^ in D1–Cre or A2A–Cre mice (Fig. 1a,b). Fluorescence changes with 473–nm and 561–nm excitation were collected by a spectrometer and spontaneous locomotion was recorded simultaneously by a frame-synchronized video camera (Fig. 1c). Linear unmixing successfully separated the fluorescence signal from gDA3m (green), Venus–iGluSnFR (amber), and sRGECO (magenta) (Fig. 1d). Nine body landmarks were tracked to calculate single-animal pose-estimation from the videos using DeepLabCut^24,25^ (Fig. 1e). Eight locomotor features were extracted for unsupervised behavior classification of spontaneous behaviors, which were then aligned to the photometry signals (Fig. 1f,h).

**Figure 1.**
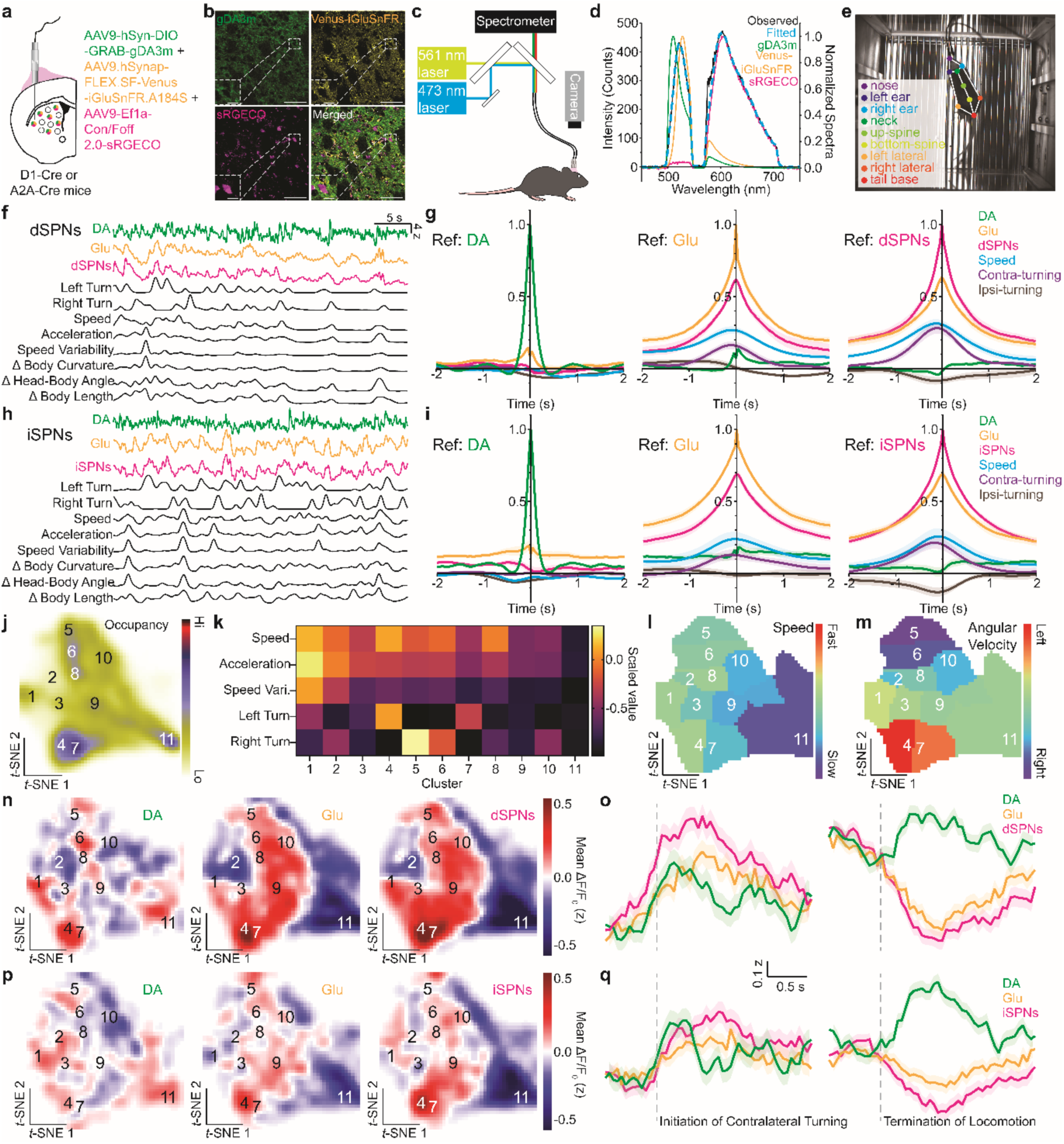
Striatal dopamine dynamics do not correlate with spontaneous movement. **a**, Viral strategy for gDA3m, Venus–iGluSnFR and sRGECO in DLS with Cre-on AAVs and optical fiber probe placement in D1–Cre or A2A–Cre mice. **b,** Confocal images showing indicator expression in dSPNs. Scale bars: 50 µm (main) and 15 µm (inset). **c,** Spectrally resolved fiber photometry system and top-view camera configuration. **d,** Example observed emission spectrum with reference spectra and linear-unmixing fit. **e,** Video frame with tracked body parts. **f,h,** Example DA, glutamate, SPN activity and movement-feature traces from D1–Cre (**f**) and A2A–Cre (**h**) mice. **g,i,** Auto- and cross-correlation summaries for DA, glutamate, SPN activity, speed and turning in D1–Cre (**g**) and A2A–Cre (**i**) mice. **j,** Smoothed occupancy map of *t*-distributed stochastic neighbor embedding (*t*-SNE) from locomotion features. **k-m,** Cluster-wise locomotion features, speed and angular velocity. **n,p,** DA, glutamate and SPN activity mapped across *t*-SNE behavior space in right-hemisphere dSPNs (**n**) and iSPNs (**p**). **o,q,** Average signals during contralateral turning and locomotion termination in dSPNs (**o**) and iSPNs (**q**). Data included in the analysis are from 7 hemispheres in 4 D1–Cre mice and 8 hemispheres in 4 A2A–Cre mice (g,i-q).

Cross-correlation analysis showed that DA was weakly coupled to ongoing spontaneous movement. At zero lag, DA-referenced correlations with glutamate, SPN activity, speed, contralateral turning, and ipsilateral turning were close to zero in both SPN recordings (dSPNs, Fig. 1g, *left*; iSPNs, Fig. 1i, *left*). By contrast, glutamate and SPN activity were strongly correlated with each other and with locomotor features, indicating that glutamatergic input and SPN activity, but not DA, were continuously aligned with spontaneous kinematics (Fig. 1g,i, *middle* and *right*).

We next asked whether DA was organized by broader locomotor states. Frame-wise movement features were embedded into a two-dimensional *t*-distributed stochastic neighbor embedding (*t*-SNE) manifold and partitioned into 11 behavioral clusters, using density-based clustering (Fig. 1j). Cluster-wise feature maps identified cluster 11 as a low-mobility state, whereas projections of speed and angular velocity separated fast locomotion and directional turning, with left and right turns enriched in distinct regions of the manifold: left turns were predominantly represented in cluster 4 and right turns were enriched in cluster 5 (Fig. 1k-m and Extended Data Fig. 1a-g). Mapping photometry signals onto this behavioral space showed that glutamate and SPN activity formed structured, cluster-consistent patterns in both dSPN and iSPN recordings. DA signals, however, were spatially heterogeneous and did not recapitulate the speed, turning or SPN activity maps (Fig. 1n,p and Extended Data Fig. 2a-f).

Cluster-triggered averages further separated DA from spontaneous movement (Extended Data Fig. 3). During contralateral turning (cluster 4 from right striatum and cluster 5 from left striatum), DA, glutamate and SPN activity increased in both pathways (Fig. 1o,q, *left*). After locomotion termination, however, DA increased while glutamate and SPN activity decreased (Fig. 1o,q, *right*). Thus, although striatal DA transients occurred around selected motor transitions^26^, DA dynamics were not tuned to spontaneous speed, turning or locomotor-state structure, unlike the coordinated glutamate and SPN activity signals.

### Peri-event DA levels predicted SPN input-output efficacy

Despite glutamate and SPN activity being strongly correlated with each other, we found the relative magnitude of peaks in Venus–iGluSnFR and sRGECO were not always consistent in dSPNs, (Fig. 2a), indicating wide variability in input-output efficacy of glutamatergic signaling in dSPNs. To probe the role of DA in this variability, we annotated events across different clusters based on paired peaks in Venus–iGluSnFR and sRGECO signals, using their rapid rising phase for temporal alignment with the gDA3m signal (Fig. 2b,c). We then grouped events into three categories based on the output/input ratio (sRGECO / Venus-iGluSnFR: *low* < 0.67 < *medium* <1.5 < *high*; Fig. 2d). The distribution across three categories was 24.21% (46/190) low-efficacy, 51.58% (98/190) medium-efficacy and 24.21% (46/190) high-efficacy (Fig. 2e). DA fluorescence (gDA3m) showed a positive peak before high-efficacy events, a negative peak before low-efficacy events, and showed no peaks in medium-efficacy events (Fig. 2d,e). Quantification confirmed that DA fluorescence (−0.2 ∼ +0.1 s) scaled with output-input efficacy (*low*: −0.6008 ± 0.1399; *medium*: −0.09176 ± 0.07039; *high*: 0.4471 ± 0.1174; Fig. 2f). We next asked whether DA could predict dSPN input-output efficacy. We sorted paired glutamate dSPN events into low-, medium-, and high-DA groups based on the peri-event gDA3m signal (Fig. 2g). The sRGECO / Venus– iGluSnFR ratio progressively increased with DA level (Fig. 2h), and quantification confirmed significantly greater efficacy in the high-DA group (Fig. 2i). Thus, higher DA predicts enhanced dSPN input-output efficacy.

**Figure 2.**
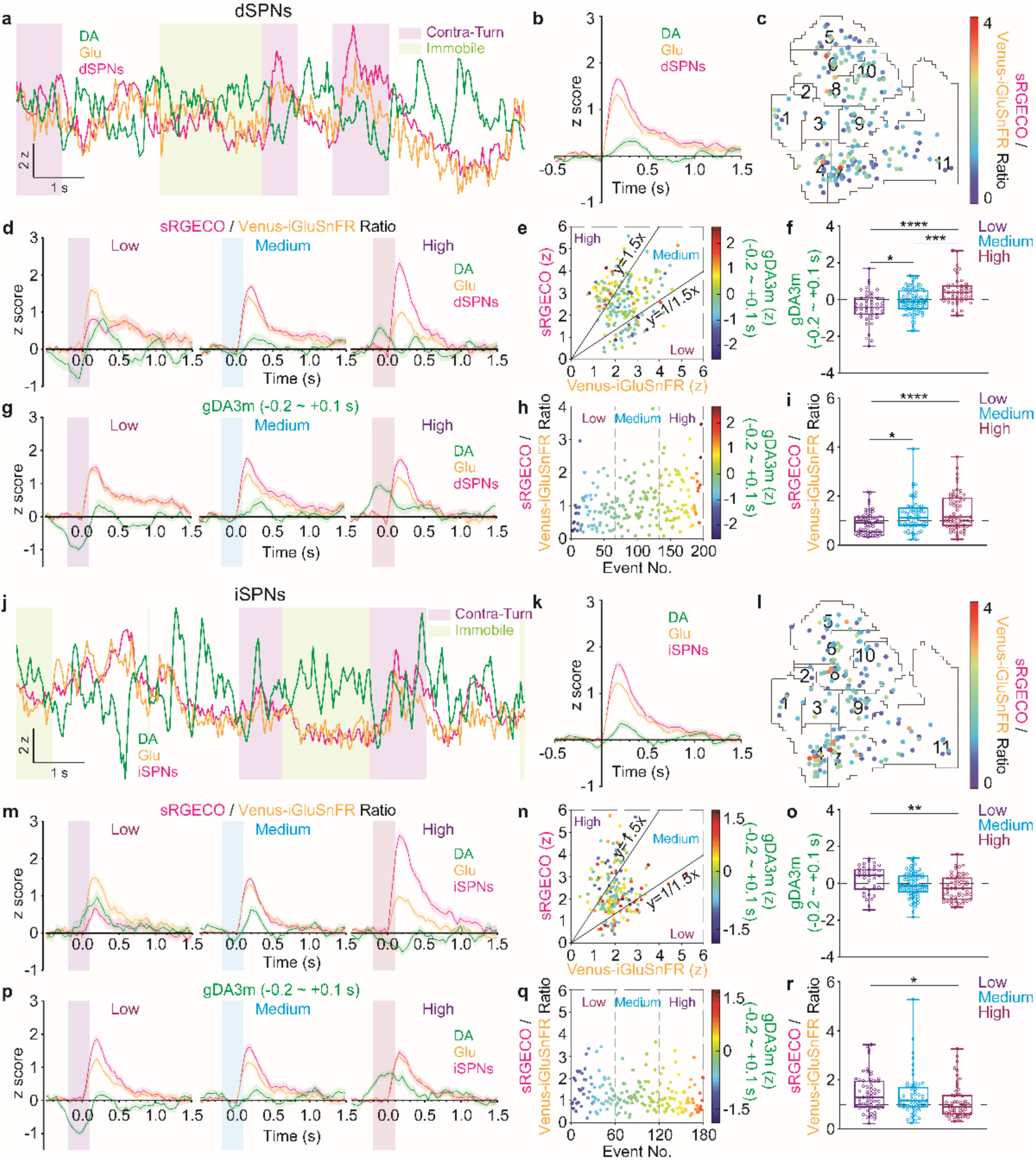
Preceding striatal dopamine tone bi-directionally modulates SPNs neural activity. **a,j**, Example of DA, glutamate, SPN activity and contra-turn or immobile clusters from D1–Cre (**a**) or A2A– Cre (**j**) mice. **b,k,** Summary of all events (represented as mean ± SEM) from dSPNs (**b**) or iSPNs (**k**). **c,l,** Event map of *t*-SNE behavior space for dSPNs (**c**) or iSPNs (**l**). The ratio of sRGECO / Venus–iGluSnFR was color coded for each event. **d,m,** Summary of the events (represented as mean ± SEM) in the *low* (*left*), *medium* (*middle*) and *high* (*right*) sRGECO / Venus–iGluSnFR ratio groups from dSPNs (**d**) or iSPNs (**m**). **e,n,** Scatterplots showing the relationship between the magnitude of Venus–iGluSnFR (x axis) and sRGECO (y axis) and gDA3m (the average Z score of 0.2 s before to 0.1 s after the transients onset, color coded) for dSPNs (**e**) or iSPNs (**n**). **f,o,** Comparison of gDA3m (the average Z score of 0.2 s before to 0.1 s after the transients onset) in dSPNs (**f**) or iSPNs (**o**). **g,p,** Summary of the events (represented as mean ± SEM) grouped by mean gDA3m (−0.2 ∼ +0.1 s) into *low* (*left*), *medium* (*middle*) and *high* (*right*) of from dSPNs (**g**) or iSPNs (**p**). **h,q,** Scatterplots showing the sRGECO / Venus–iGluSnFR ratio (y axis) across events sorted by mean gDA3m (−0.2 ∼ +0.1 s; x axis: event number), with mean gDA3m color coded, for dSPNs (**h**) or iSPNs (**q**). **i,r,** Comparison of the ratio of sRGECO / Venus–iGluSnFR from dSPNs (**i**) or iSPNs (**r**). Events included in the analysis are 190 events from 7 hemispheres in 4 D1–Cre mice (**b-i**); 175 events from 8 hemispheres in 4 A2A–Cre mice (**k-r**). *, p < 0.05; **, p < 0.01; ***, p < 0.001; ****, p < 0.0001, one-way ANOVA followed by Tukey’s multiple comparisons test in **f,i,o,r**.

We also applied the same analysis to probe the effects of DA on iSPNs (Fig. 2j). Paired peaks in Venus–iGluSnFR and sRGECO were annotated from recordings in A2A–Cre mice (Fig. 2k,l). Events were grouped by input-output efficacy (low 20.57% [36/175], medium 48.57% [85/175], high 30.85% [54/175], Fig. 2m), revealing a negative relationship between peri-event DA (−0.2 ∼ +0.1 s) and iSPN efficacy (Fig. 2n,o). When events were grouped by DA level (Fig. 2p), the iSPN response relative to glutamatergic input decreased as DA increased (Fig. 2q), and the high-DA group showed significantly lower efficacy. (Fig. 2r). These results suggest that DA level positively correlates with event-based input-output efficacy in dSPNs and negatively correlates with iSPNs.

### Effect of dopamine reuptake blockade on input-output efficacy and locomotion

To further validate dopaminergic modulation of striatal SPNs and locomotion, we assessed the effects of DA reuptake blockade on neural signals and behavior. Mice were administered methylphenidate (MPH, 10 mg kg^-1^) immediately after a 5-min baseline recording (Pre-MPH), followed by a second 5-min recording (Post-MPH) 30 min later (gDA3m traces shown in Fig. 3a,k). Peaks in gDA3m fluorescence were detected to quantify MPH-induced DA dynamic changes: in both dSPNs and iSPNs (Fig. 3b,l). MPH increased the amplitude (Fig. 3c,m), frequency (Fig. 3d,n), and area under the curve (Fig. 3e,o) of DA transients and prolonged their decay kinetics (Fig. 3f,p).

**Figure 3.**
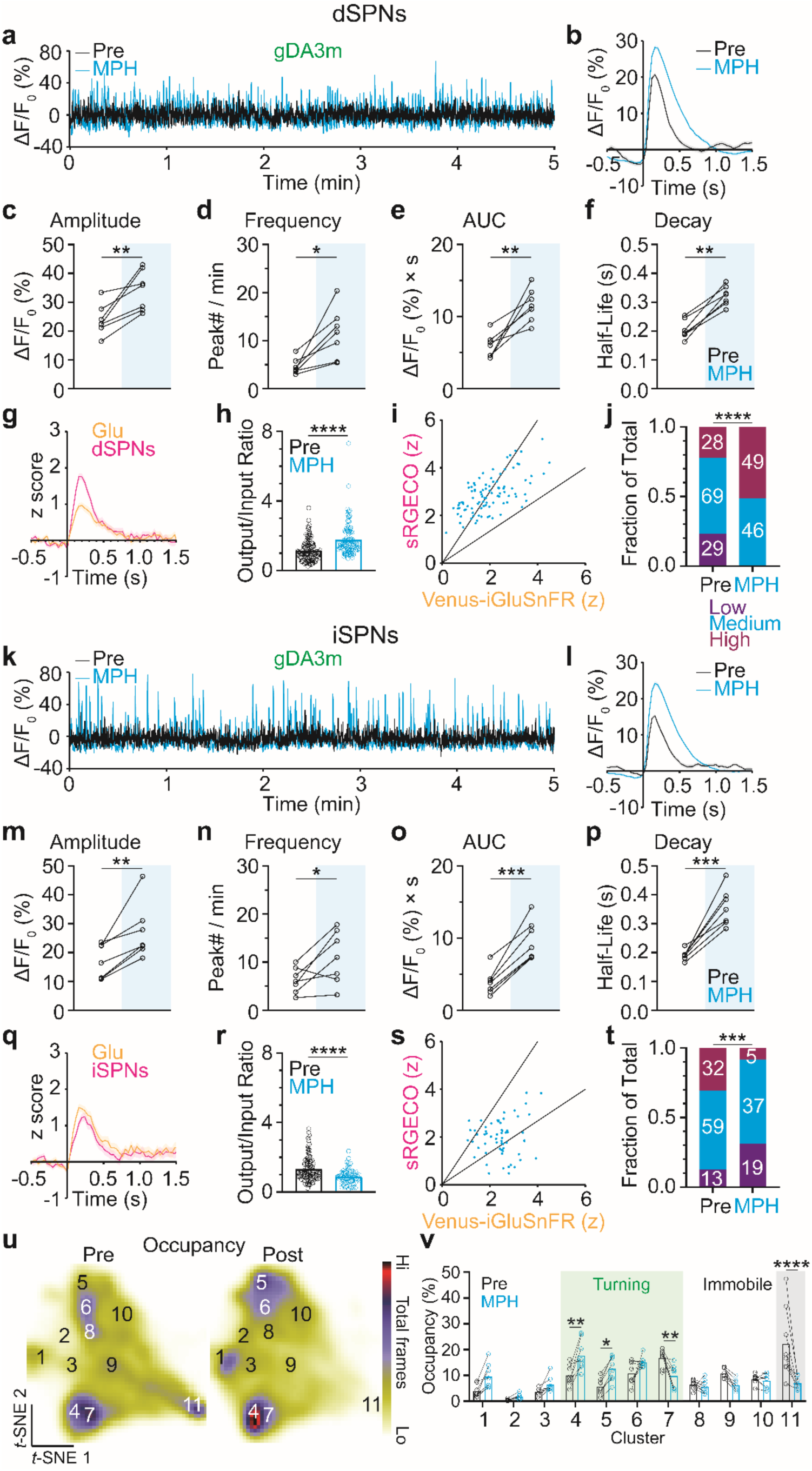
Dopamine reuptake blockade reshapes input-output efficacy to increase mobility. **a,k**, Representative fluorescence traces of dopamine before (black) and after administration of MPH (blue) from dSPNs (**a**) or iSPNs (**k**). **b,l,** Summary of gDA3m peaks (represented as mean ± SEM) before (black) and after administration of MPH (blue) from dSPNs (**b**) or iSPNs (**l**). **c,m,** Comparison of peak amplitude of gDA3m before (black) and after MPH (blue) from dSPNs (**c**) or iSPNs (**m**). **d,n,** Comparison of peak frequency of gDA3m before (black) and after MPH (blue) from dSPNs (**d**) or iSPNs (**n**). **e,o,** Comparison of peak AUC of gDA3m before (black) and after MPH (blue) from dSPNs (**e**) or iSPNs (**o**). **f,p,** Comparison of decay kinetics of gDA3m before (black) and after MPH (blue) from dSPNs (**f**) or iSPNs (**p**). **g,q,** Summary of all events (represented as mean ± SEM) from dSPNs (**g**) or iSPNs (**q)** after MPH. **h,r,** Comparison of output/input ratio before (black) and after MPH (blue) from dSPNs (**h**) or iSPNs (**r**). **i,s,** Scatterplots showing the relationship between the magnitude of Venus–iGluSnFR and sRGECO from dSPNs (**i**) or iSPNs (**s)** after MPH. **j,t,** Distribution of events across *low*, *medium* and *high* output/input ratio groups before and after MPH from dSPNs (**j**) or iSPNs (**t**). **u,** Smoothed histogram of *t*-SNE embeddings from before (Pre, left) and after MPH (Post, right). **v,** Comparison of occupancy changes in clusters before (black) and after MPH (blue). Data included in the analysis are from 7 hemispheres in 4 D1–Cre mice and 7 hemispheres in 4 A2A–Cre mice. Events included in the analysis are as shown in **j** for **g-i**, and **t** for **q-s**. *, p < 0.05; **, p < 0.01; ***, p < 0.001, paired *t* test in **c-f, m-p**. ****, p < 0.0001, unpaired *t* test in **h,r**. ***, p < 0.001; ****, p < 0.0001, Fisher’s exact test in **j,t**. *, p < 0.05; **, p < 0.01; ****, p < 0.0001, one-way ANOVA followed by Tukey’s multiple comparisons test in **v**.

Event-based analysis of Venus–iGluSnFR and sRGECO showed that DA reuptake blockade increased the ratio of sRGECO / Venus–iGluSnFR in dSPNs (Fig. 3g,h) whereas it decreased this measure in iSPNs (Fig. 3q,r), consistent with opposite changes in Venus–iGluSnFR and sRGECO (Extended Data Fig. 5a-d). Furthermore, the distribution across efficacy categories shifted toward high efficacy in dSPNs (Fig. 3i,j) and toward low efficacy in iSPNs (Fig. 3s,t).

Unsupervised behavior quantification revealed a significant redistribution of cluster occupancy after MPH (Fig. 3u): mice spent more time in large left turns (cluster 4) and large right turns (cluster 5), whereas they spent correspondingly less time in immobile behavior (cluster 11, Fig. 3v). These results suggest that DA reuptake blockade with MPH increases input-output efficacy of dSPNs and decreases it in iSPNs, concomitant with a redistribution of locomotor-state occupancy.

### Effects of dopamine depletion on input-output efficacy and locomotion

To further evaluate dopaminergic modulation of striatal SPNs and locomotion, we next examined the effects of DA depletion on neural signals and behavior. Mice underwent a 5-min baseline recording (Pre-RES) prior to reserpine administration (RES; 2.5 mg kg^−1^), followed by a second 5-min recording (Post-RES) 6 h after injection (gDA3m traces shown in Fig. 4a,k). In both dSPNs and iSPNs, RES markedly reduced the amplitude (Fig. 4c,m), frequency (Fig. 4d,n), and area under the curve (Fig. 4e,o) of DA transients and accelerated their decay kinetics (Fig. 4f,p).

**Figure 4.**
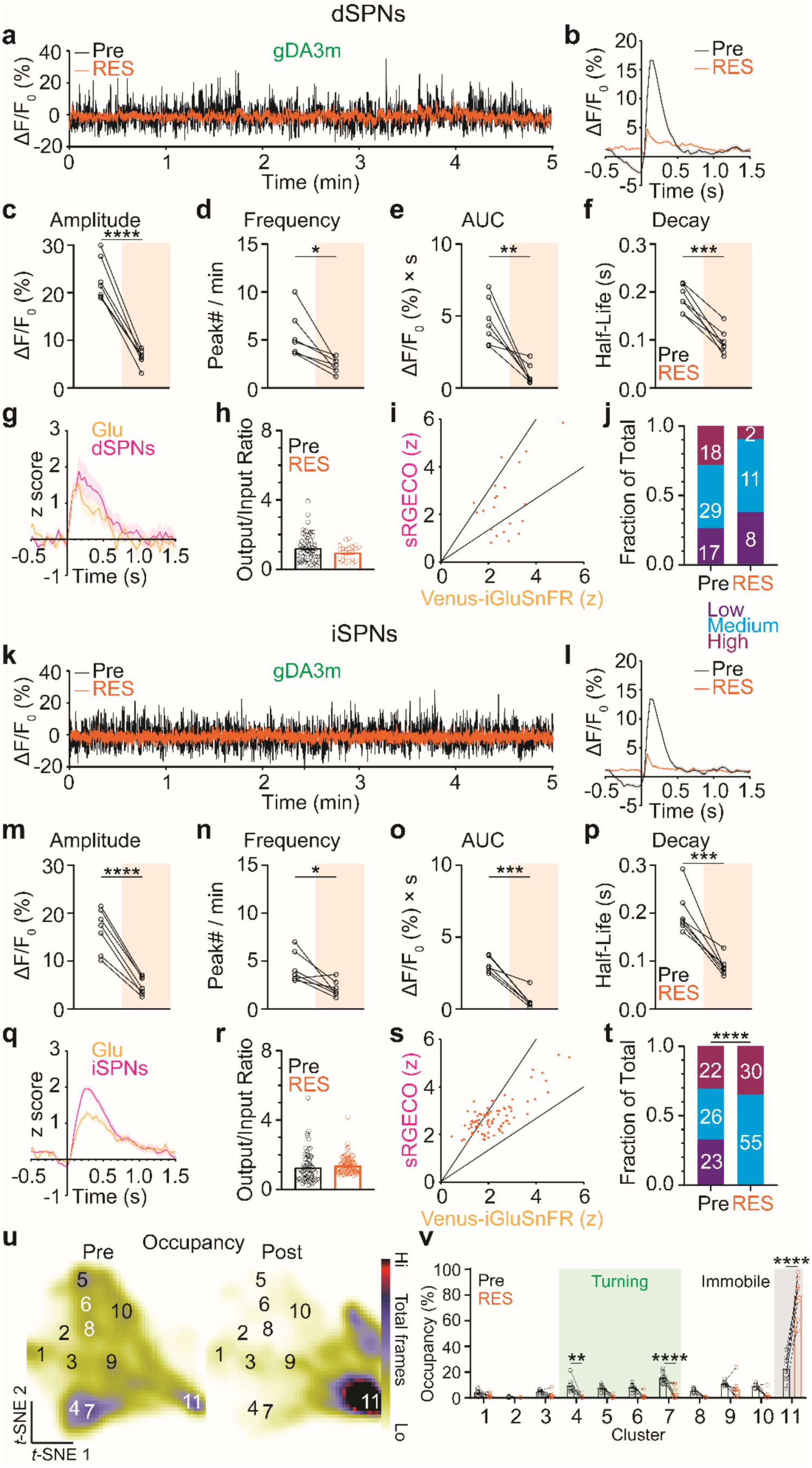
Dopamine depletion reshapes input-output efficacy to decrease mobility. **a,k**, Representative fluorescence traces of dopamine before (black) and after administration of RES (orange) from dSPNs (**a**) or iSPNs (**k**). **b,l,** Summary of gDA3m peaks (represented as mean ± SEM) before (black) and after administration of RES (orange) from dSPNs (**b**) or iSPNs (**l**). **c,m,** Comparison of peak amplitude of gDA3m before (black) and after RES (orange) from dSPNs (**c**) or iSPNs (**m**). **d,n,** Comparison of peak frequency of gDA3m before (black) and after RES (orange) from dSPNs (**d**) or iSPNs (**n**). **e,o,** Comparison of peak AUC of gDA3m before (black) and after RES (orange) from dSPNs (**e**) or iSPNs (**o**). **f,p,** Comparison of decay kinetics of gDA3m before (black) and after RES (orange) from dSPNs (**f**) or iSPNs (**p**). **g,q,** Summary of all events (represented as mean ± SEM) from dSPNs (**g**) or iSPNs (**q)** after RES. **h,r,** Comparison of output/input ratio before (black) and after RES (orange) from dSPNs (**h**) or iSPNs (**r**). **i,s,** Scatterplots showing the relationship between the magnitude of Venus–iGluSnFR and sRGECO from dSPNs (**i**) or iSPNs (**s)** after RES. **j,t,** Distribution of events across *low*, *medium* and *high* output/input ratio groups before and after RES from dSPNs (**j**) or iSPNs (**t**). **u,** Smoothed histogram of *t*-SNE embeddings from before (Pre, left) and after RES (Post, right). **v,** Comparison of occupancy changes in clusters before (black) and after RES (orange). Data included in the analysis are from 7 hemispheres in 4 D1–Cre mice and 7 hemispheres in 4 A2A–Cre mice. Events included in the analysis are as shown in **j** for **g-i**, and **t** for **q-s**. *, p < 0.05; **, p < 0.01; ***, p < 0.001; ****, p < 0.0001, paired *t* test in **c-f, m-p**. n.s., p > 0.05, unpaired *t* test in **h,r**. n.s., p > 0.05; ****, p < 0.0001, Fisher’s exact test in **j,t**. **, p < 0.01; ****, p < 0.0001, one-way ANOVA followed by Tukey’s multiple comparisons test in **v**.

Event-based analysis of Venus–iGluSnFR and sRGECO further indicated that DA depletion produced no significant change in the output/input ratio of glutamatergic signaling in dSPNs (Fig. 4g,h) and iSPNs (Fig. 4q,r), accompanied by increased Venus–iGluSnFR signals in dSPNs and increased sRGECO signals in iSPNs (Extended Data Fig. 6a-d). In contrast to MPH, the distribution across efficacy categories weakly shifted toward low efficacy in dSPNs (Fig. 4i,j) and toward higher-efficacy categories in iSPNs (Fig. 4s,t).

Unsupervised behavioral quantification revealed a pronounced redistribution of cluster occupancy following RES (Fig. 4u). In contrast to MPH, mice spent more time in the immobile state (cluster 11) and correspondingly less time in turning clusters (clusters 4 and 7, Fig. 4v). These results suggest that DA depletion with RES is accompanied by pathway-specific changes in SPN input-output efficacy and a redistribution of locomotor-state occupancy toward immobility.

## Discussion

In this study, we combined genetically encoded fluorescent indicators with triple-color spectrally resolved fiber photometry to simultaneously measure DA, glutamate release and SPN activity in the DLS during mouse locomotion. We found that DA dynamics did not track with spontaneous locomotor kinematics or states. Instead, DA transients preceding coupled increases in glutamate release and SPN activity predicted SPN input-output efficacy with opposite relationships across pathways: higher DA was associated with greater input-output efficacy of dSPNs and lower efficacy in iSPNs, providing *in vivo* support for the classical model in which D1 receptor signaling promotes direct-pathway excitability and D2 receptor signaling suppresses indirect-pathway excitability^27^. These conclusions were further supported by pharmacological manipulations that elevation or depletion of extracellular DA produced corresponding, pathway-specific changes in SPN input-output efficacy and reorganized locomotor-state occupancy.

Unsupervised behavior classification^28,29^ revealed locomotion-linked neural signatures in DLS. The observation that Venus–iGluSnFR and sRGECO increased in both dSPNs and iSPNs during directional movement (cluster 4 and 7, Fig. 1n-q; Extended Data Figs. 2b,c,e,f and 3) is consistent with prior findings that concurrent activation of both SPN populations within one hemisphere precedes the initiation of contraversive movements^30,31^. Surprisingly, DA signals also showed increases following the termination of locomotion (cluster 11). This finding is consistent with our earlier observation that correlations between DA and speed or turning were near zero in both SPN populations, further supporting the conclusion that DA was weakly coupled to ongoing spontaneous movement.

We found that DA correlates with the event-based output/input ratio of dSPNs and iSPNs in opposite directions (Fig. 2f,i,o,r). This result is consistent with the prevailing receptor-signaling framework, in which D1R signaling increases cAMP production and PKA activity to enhance dSPN excitability, whereas D2R signaling suppresses cAMP/PKA signaling to reduce iSPN excitability^2–4,9–13^. Notably, the smaller separation among efficacy categories in iSPNs (Fig. 2o,r) relative to dSPNs (Fig. 2f,i) aligns with a stronger dopaminergic gain on dSPNs in modulating excitability. It’s also worth noting that DA signals were not linearly related to neural activity or glutamate release during spontaneous locomotion (Extended Data Fig. 4), suggesting that moment-to-moment SPN activity is not governed by a simple linear relationship with either signal; instead, the contributions of DA and glutamate may be partially dissociable and dependent on pathway, behavioral state, and timescale^14^. In addition, we observed that DA efficacy coupling was temporally constrained to a narrow window around glutamate- and activity-associated transients, within ∼200 ms before and ∼100 ms after the onset of Venus–iGluSnFR and sRGECO peaks, consistent with the expected kinetics of GPCR-dependent cascades, which typically evolve over hundreds of milliseconds^32^.

Consistent with the correlational findings, bidirectional pharmacological manipulation of extracellular DA produced opposing effects on SPN input-output efficacy and locomotor-state occupancy. DA reuptake blockade^33^ increased both the event-based output/input ratio and the fraction of high-efficacy events in dSPNs, while reducing these measures in iSPNs. Conversely, DA depletion^34^ shifted the efficacy distribution in dSPNs toward lower categories, and in iSPNs toward higher categories. Larger shifts in input-output efficacy were observed in dSPNs following MPH administration (Fig. 3j) and in iSPNs following RES administration (Fig. 4t). This pathway-specific asymmetry may partly reflect differences in the basal activation of D1 and D2 receptors, potentially related to the greater apparent affinity of D2 receptors for DA^35^. At the behavioral level, DA reuptake blockade redistributed occupancy away from the immobile state (cluster 11, p < 0.0001) and the smaller turning state (cluster 7, p < 0.01) toward larger turning states (cluster 4, p < 0.01; cluster 5, p < 0.05), whereas DA depletion produced a broader reorganization, with the immobile state (cluster 11, p < 0.0001) becoming the dominant state and turning states such as cluster 4 (p < 0.01) and 7 (p < 0.0001) strongly reduced. Together, these findings suggest that dopaminergic tone not only shifts pathway-specific input-output efficacy but can also redistribute the behavioral state space through which animals’ express locomotion.

Our study demonstrated that *in vivo* SPN activity is tightly synchronized with glutamate release, while the relative scaling between input and output varies substantially across time and behavioral state. Our results indicate that DA accounts for part of this variability by dynamically tuning the effective input-output gain of glutamatergic transmission in a receptor- and pathway-dependent manner.

In summary, our results support a model in which DA dynamics tune pathway-specific SPN input-output efficacy in the DLS, thereby reorganizing locomotion-related behavioral signatures. More broadly, our approach provides an *in vivo* window into how neuromodulators reshape neurotransmitter-driven signaling while explicitly incorporating behavioral-state structure. This framework could be extended in future studies to test how neuromodulator-dependent gain regulation interacts with glutamatergic inputs during motor learning and goal-directed behaviors.

## Methods

### Mice

All animal protocols were approved by the National Institute of Environmental Health Sciences Animal Care and Use Committee. Experiments were carried out using 2 to 6-month-old male and female mice. D1–Cre (MMRRC_036916-UCD) and A2A–Cre (MMRRC_036158-UCD) mice were obtained from the Mutant Mouse Resource Research Centers (MMRRC). All mice were housed under reverse light cycle conditions and had access to food and water *ad libitum* at 18–23 °C with 40–60% humidity. Mice were group housed before optical probe implantation surgery and were singly housed after surgery.

### Viral vectors

pAAV-hSyn-DIO-GRAB-gDA3m (a.k.a gDA3m, Addgene plasmid# 208707; http://n2t.net/addgene:208707; RRID:Addgene_208707) was a gift from Dr. Yulong Li. pAAV.hSynap-FLEX.SF-Venus-iGluSnFR.A184S (a.k.a Venus-iGluSnFR, Addgene plasmid# 106183; http://n2t.net/addgene:106183; RRID:Addgene_106183) was a gift from Dr. Loren Looger. pAAV-Ef1a-Con/Foff 2.0-sRGECO was a gift from Dr. Karl Deisseroth & INTRSECT 2.0 Project (a.k.a sRGECO, Addgene plasmid# 137127; http://n2t.net/addgene:137127; RRID:Addgene_137127).

AAV9-DIO-Venus–iGluSnFR and AAV9-ConFoff-sRGECO were recovered in house by the NIEHS Viral Vector Core and had titers of 1.5 to 2.0 × 10^13^ genome copies per ml with AAV9 capsid. AAV9-DIO-gDA3m was obtained from Biohippo, Inc. and had titers of 3.74 × 10^12^ genome copies per ml.

### Stereotaxic microinjections of AAVs

A mixture of AAVs containing gDA3m, Venus-iGluSnFR, sRGECO at a 1:2:2 volume ratio was injected at 500 nl per site at a rate of 100 nl per min through a Hamilton Neuros syringe with a 30–gauge needle via standard stereotaxic procedures with D1–Cre or A2A–Cre mice under isoflurane anesthesia^36^. The needle was left in place for ten more minutes before withdrawal. The coordinates used for targeting the dorsolateral striatum (DLS) were AP +0.50 mm, ML ±2.20 mm from Bregma, and DV −3.00 mm from the brain surface.

### Fiber photometry-guided surgery for optical fiber probe implantation

Three weeks after the virus injection, mice were subjected to a stereotaxic surgery for the implantation of the optical fiber probe as previously described^31^. Briefly, holes were drilled bilaterally through the skull above the DLS (AP: +0.5 mm, ML: ±2.4 mm from Bregma), using a #1/2 (0.027’’ diameter) drill bit. Three more bur holes were drilled in the skull for anchoring screws. Then, the spectrally resolved fiber photometry system was powered on^20,31^. The output power of the 473 nm and 561 nm lasers was set at 50-75 µW, measured at the end of the patch cable. Next, the optical fiber probe was connected to the patch cable. In the spectrometer operation software OceanView (Ocean Optics), the background spectrum detected by the spectrometer (QE-Pro, Ocean Optics) was subtracted when the room light was turned off. The optical fiber probe was lowered above the brain surface through the burr hole, then further lowered until the emission spectrum of Cre-dependent fluorescent proteins started to appear in OceanView. The probe was then lowered slowly while the magnitude of the emission spectrum was being monitored until it reached a plateau. The final probe tip location in the DLS was approximately 2.1-2.2 mm below the brain surface. The probe was then fixed in place with a generous amount of dental acrylic (Jet, Lang Dental Mfg. Co.). The mice were allowed one week to recover before experiments proceeded.

### Spectrally resolved fiber photometry

The fiber photometry recordings were carried out in freely moving mice in an open-top mouse operant chamber (21.6 cm × 17.8 cm × 12.7 cm, ENV-307W-CT, Med Associates, Inc) housed in a sound attenuating box (ENV-017M, Med Associates, Inc). Fluorescence spectra were acquired by a spectrometer as described previously^31^ using a 19 ms integration time, triggered by 25 Hz TTL pulses sent from a digital output module (DIG-726TTL, Med-Associates) on a customized mouse operant conditioning package (Med Associates, Inc). The output power of the 473 nm and 561 nm lasers measured at the end of the patch cable was set at about 50-75 µW. A digital video camera (Grasshopper3 GS3-U3-23S6M-C, FLIR Integrated Imaging Solutions), triggered frame by frame by the same TTL pulses as the spectrometer, was used to record the animal’s behavior. Spectral linear unmixing was performed on the acquired fluorescence emission spectra to isolate individual spectral components using a customized program written in R (see code availability statement below). To correct the fluorescence fading, we first applied linear regression or single decay non-linear regression to fit the unmixed coefficients plotted over time and then used the fitted curve as the theoretical baseline (F_0_) to calculate ΔF/F_0_%. Z scores were calculated by z = (x - m) / σ, where x is the ΔF/F_0_% at a given time, m is the mean of ΔF/F_0_% values in a single recording session, and σ is the standard deviation of all ΔF/F_0_% values in the same recording session. The detection threshold for a fluorescence transient was defined as µ + 3σ, where µ and σ were the mean and the standard deviation of the 500-ms fluorescence baseline period preceding the interrogated data point.

### Pharmacology recording with dopamine reuptake blockade and dopamine depletion

Methylphenidate (Sigma-Aldrich, M2891-100MG) was first dissolved in sterile 0.9% saline to make a 10 mg ml^-1^ stock solution. Then, 100 µl of stock solution was diluted with 900 µl of sterile 0.9% saline to make the final injection solution. The mice were recorded for 5 min before methylphenidate (10 mg kg^-1^) was injected intraperitoneally (Pre-MPH recording). A second 5-min recording was repeated at 30 min after methylphenidate injection (Post-MPH recording).

Dopamine depletion recordings were performed 48 hours after dopamine reuptake blockade recordings. Reserpine (Sigma-Aldrich, R0875-1G) was first dissolved in DMSO to make a 5 mg ml^-1^ stock solution. Then, 50 µl of stock solution was dissolved with 950 µl of sterile 0.9% saline containing 0.02% ascorbic acid to make the final injection solution. The mice were recorded for 5 min before reserpine (2.5 mg kg^-1^) was injected intraperitoneally (Pre-RES recording). A second 5-min recording was repeated at 6 h after reserpine injection (Post-RES recording).

Mice were perfused immediately after Post-RES recording.

### Behavior tracking and movement analysis

The animal’s behavior was captured by a digital video camera (Grasshopper3 GS3-U3-23S6M-C, FLIR Integrated Imaging Solutions, Inc.), which was triggered by the same TTL pulses that triggered the spectrometer. Recorded videos were analyzed with DeepLabCut v2.2.3^24,25^ in Anaconda to measure the coordinates of “nose”, “left ear”, “right ear”, “neck”, “up-spine”, “bottom-spine”, “left-lateral”, “right-lateral” and “tail base”.

### Feature definition

Eight per-frame behavioral features were computed from the DeepLabCut coordinates using POSEIDON, a custom Python package for unsupervised behavioral embedding. All features were computed at a frame rate of 25 Hz.

1. Velocity (px s⁻¹): the Euclidean displacement of the “body centroid”, defined as the mean of “tail base”, “nose”, “left ear”, and “right ear” coordinates, between consecutive frames, multiplied by the frame rate.
2. (2–3) Left and right windowed turn rate (rad s⁻¹): the per-frame heading change, computed as the wrapped angular difference of the “body centroid” to “nose” vector between successive frames, was summed over a centered 0.5–second sliding window (13 frames) and divided by the window length. The resulting net turn rate was then split into two non-negative channels, left (counterclockwise) and right (clockwise), to prevent cancellation in cluster averages.
3. Spine curvature change (rad/s): the sum of the angles at two spine joints, computed as the angle between the vector from “tail base” to “bottom spine” and the vector from “bottom spine” to “upper spine” plus the angle between the vector from “bottom spine” to “upper spine” and the vector from “upper spine” to “nose”, where each angle is the arccosine of the normalized dot product of the two vectors. The absolute difference between consecutive frames is multiplied by the frame rate and averaged over the analysis window (0.5 s). Larger values indicate more rapid changes in spinal curvature, irrespective of the direction of bending.
4. Head-body angle change (rad/s): the angle between the the vector from “tail base” to “neck” and the vector from “neck” to “nose”, computed via the same arccosine method. The absolute difference between consecutive frames is multiplied by the frame rate and averaged over the analysis window (0.5 s). Larger values indicate more rapid changes in head orientation relative to the body, irrespective of direction.
5. Spine length change (px/s): the summed Euclidean distances of “tail base” to “bottom spine”, “bottom spine” to “upper spine”, and “upper spine” to “neck”. The absolute difference between consecutive frames is multiplied by the frame rate and averaged over the analysis window (0.5 s). Larger values indicate more rapid extension or contraction of the body axis.
6. Acceleration (px s⁻²): the absolute per-frame change in velocity, averaged over a centered 0.5–second rolling window.
7. Speed variability (px² s⁻²): the variance of velocity computed over the same 0.5 second rolling window.

### Feature preprocessing

Before embedding, the eight features were preprocessed in four steps. First, each feature was globally clipped to its 1st and 99th percentiles across all sessions to reduce the influence of outliers. Second, within each session, each feature was temporally smoothed with a one-dimensional Gaussian filter (σ = 6.25 frames, equivalent to 0.25 s at 25 Hz). Third, any remaining NaN values were filled by forward fill, backward fill, and finally zero fill. Fourth, each feature was independently rescaled to the range [−1, 1] within each session using min-max normalization. Session-wise scaling ensured that animals with different baseline movement speeds contributed equally to the embedding.

### Unsupervised behavior embedding and clustering

Unsupervised behavioral classification was performed using a two-round *t*-SNE embedding procedure with importance sampling, adapted from previously described methods^28^. In round one, approximately 12,000 frames were sampled uniformly across all recording sessions with equal allocation per session without replacement. The sampled feature vectors, each with eight dimensions after preprocessing, were embedded into two dimensions using t-distributed stochastic neighbor embedding (*t*-SNE; perplexity = 100, random initialization). A multilayer perceptron (MLP) regressor (hidden layer sizes 400, 200, 50; maximum 1,000 training iterations) was then trained to approximate the mapping from the 8-dimensional feature space to the 2D *t*-SNE coordinates. This MLP was used to project all frames from every session into the same 2D embedding space, allowing for consistent comparison across animals and conditions.

The projected points were binned into a 50 by 50 two-dimensional histogram and smoothed with a 2D Gaussian filter (σ = 2.5 bins). Local density maxima were identified, excluding border bins. Bins below the 30th percentile of the density distribution were masked. Watershed segmentation was then applied to the inverted density map, seeded by the labeled local maxima, to define coarse behavioral clusters. Each frame was assigned to the cluster corresponding to its histogram bin in the watershed label map.

To improve the resolution of behaviorally relevant clusters, a second embedding round was performed with importance sampling. The round 1 clusters whose frames had the highest overlap with the top 20th percentile of the left and right windowed turn rate features were identified as turning related clusters. A new training set of approximately 10,000 frames was then drawn, oversampling the identified turning clusters at a 5:2 ratio relative to all other clusters. A new *t*-SNE embedding and MLP regressor were fitted on this importance-weighted sample using the same hyperparameters as round one. All frames were re-projected into the Round 2 embedding space via the new MLP.

Final clusters were obtained by repeating the watershed procedure on the Round 2 embedding density (50 by 50 bins, Gaussian smoothing σ = 1.5 bins, 30th percentile density cutoff). Each frame across all sessions was assigned a final cluster label by binning its Round 2 embedding coordinates into the watershed labeled histogram grid.

### Neural activity in behavioral embedding space

To visualize how neural signals varied across the behavioral landscape, fiber photometry data were mapped onto the *t*-SNE embedding space. Because the spectrometer and the video camera were triggered by the same 25 Hz TTL pulse train, photometry samples were aligned one to one with behavior frames within each session with no temporal shift. For each session, every behavior frame’s 2D embedding coordinate was assigned to a bin in the same 50 by 50 histogram grid used for clustering. The mean photometry z-score value was computed for each occupied bin. Per-session maps were then averaged across sessions and smoothed with a 2D Gaussian filter. Separate maps were generated for gDA3m (DA), Venus–iGluSnFR (Glu) and sRGECO (RG) channels.

### Peri-event time histograms

Cluster-onset triggered neural responses were computed to examine how fiber photometry signals changed at the transition into specific behavioral clusters. For each cluster, onset events were defined as the first frame of a run of at least 3 consecutive frames assigned to that cluster. Peri-event time windows spanning from 0.5 seconds before to 1.5 seconds after each onset were extracted from the photometry traces. The mean ± SEM across all onset events was computed for each channel and cluster.

### Fixed brain slice preparation and imaging

Mice were transcardially perfused with 20 ml ice-cold phosphate buffered saline (PBS) immediately followed by 20 ml ice-cold 4% paraformaldehyde (PFA). Brains were post-fixed in 4% PFA overnight and then transferred to 30% sucrose in PBS for storage at 4 °C until further processing. Coronal slices were sectioned on a microtome (KS34, Thermo Fisher Scientific) at a thickness of 35 µm. Slices were mounted on slides and spectrally imaged on a Zeiss laser-scanning inverted confocal microscope (LSM 980) equipped with a multiline argon laser (458, 488 and 514nm) and two HeNe lasers (594 and 633nm) using an oil-immersion 40× objective (numerical aperture 1.3). The images were acquired and linearly unmixed using Zen 2012 Black software (Carl Zeiss).

### Annotation of events on Glu and RG channels

Potential Venus–iGluSnFR and sRGECO transients were first identified using deliberately permissive screening procedures. Glu and RG were independently smoothed using a five-sample, second-order Savitzky-Golay filter. A joint screening signal was defined as

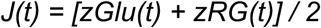

A candidate was generated when the maximum one-frame rise in *J(t)* was at least 0.86 robust standard-deviation units per 40-ms frame and the maximum joint signal between +0.04 and +0.44 s was at least 1.65. Candidates separated by less than 0.60 s were initially merged, retaining the candidate with the larger joint peak. This produced 18,201 candidates for review.

Every candidate was inspected in a graphical review interface. This interface displayed only Glu and RG and used randomized event and recording identifiers. DA traces, PRE/POST identity, pharmacological condition, genotype, animal identity, recording side and original filenames were hidden. The reviewer classified candidates as accepted, rejected or uncertain. A candidate was accepted when the waveform was judged to contain a discrete Glu/RG transient with an identifiable onset and peak and to represent a distinct event rather than baseline fluctuation, noise or a duplicate representation of a neighboring event. No single amplitude threshold was used to define acceptance. Peak amplitude was calculated relative to the local baseline, defined as the median of the preceding 2-s interval.

A frozen model for reproducibility purposes was generated after annotation. The model feature vector contained 115 Glu/RG variables: 21 Glu and 21 RG robust-z waveform samples from −0.20 to +0.60 s; 16 Glu and 16 RG one-frame derivatives; and 41 engineered morphology and interchannel features. The engineered variables included baseline scale and scale ratio; baseline slope; Glu and RG peak amplitude from 0 to +0.28 s; peak amplitude from 0 to +0.48 s; peak latency; minimum and mean Glu/RG peak; absolute peak imbalance; one-frame and two-frame rise; rise latency; Glu/RG rise imbalance; peak-to-post-event decay; Glu/RG waveform correlation; and Glu/RG derivative correlation. DA, genotype, treatment, PRE/POST status, D/A group, animal, side, recording filename and absolute recording time were excluded from the model inputs. A database-specific Extra-Trees classifier was trained to reproduce the 627 finalized event timestamps from the archived photometry recordings. The model consisted of 32 decision trees and used the Gini impurity criterion, square-root feature sampling at each split, a minimum split size of two samples, a minimum leaf size of one sample, no bootstrap resampling, and class weights balanced according to class frequency. The random seed was fixed at 20260811 to ensure deterministic model fitting. The model was implemented in Python 3.13.5 using NumPy 2.3.5, pandas 2.2.3, SciPy 1.17.0, scikit-learn 1.8.0, and joblib 1.5.3.

### DA peak detection

Positive DA peaks were detected separately in each 5-min trace. A trace-specific global baseline was estimated using the median, and robust noise was estimated using the normalized median absolute deviation. Peaks were required to exceed the global baseline by at least 3 robust standard deviations and to be separated by at least 12 samples (approximately 0.5 s). Peak width was measured at 50% of peak prominence as width at half prominence for decay kinetics. A local baseline was calculated as the median of the 2-s interval immediately preceding each event. Peak amplitude was calculated relative to the local baselines, and event AUC was calculated between the half-prominence boundaries using trapezoidal integration.

## Statistical analyses

One-way ANOVA followed by Tukey’s multiple comparisons test, two-tailed unpaired and paired *t*-tests and Fisher’s exact test were carried out using GraphPad Prism 10 (GraphPad Software). Results of the statistical analyses, including Mean ± SEM, sample sizes and P values are indicated in either the text or figure legends. Age-matched and sex-matched mice were randomly assigned to groups for all experiments. No statistical methods were used to predetermine sample sizes. Instead, group sizes were selected based on similar studies. Data collection was conducted in a blinded manner. No data were excluded from the analyses. All mice that received viral injections were assessed for viral expression and fiber tract placement. For the representative immunofluorescence images shown in the figures, the staining experiments were performed on at least three mice from each group.

## Data availability

All data are available from the corresponding authors upon reasonable request.

## Code availability

The linear spectral unmixing algorithm used in this study can be downloaded at https://www.niehs.nih.gov/research/atniehs/labs/ln/pi/iv/tools.

## Acknowledgements

We thank the NIEHS Viral Vector Core for packaging AAVs, Jeff Tucker, Erica Scappini and Rob Wine of NIEHS Fluorescence Microscopy and Imaging Center for their assistance with imaging of brain sections, Lois Wyrick and Paul Cacioppo of NIEHS Office of Communications and Public Liaison for assistance with the behavioral schematic diagram, Jicheng Li and Chengbo Meng for helpful discussions. We thank Dr. Yulong Li and Dr. Loren Looger for providing viruses for this study. This work is supported by the Intramural Research Program of the National Institutes of Health, NIH/NIEHS, of the United States (ZIA ES103310 to G.C.; ZIA ES103403 to J.E.L.). The contributions of the NIH author(s) were made as part of their official duties as NIH federal employees, are in compliance with agency policy requirements, and are considered Works of the United States Government. However, the findings and conclusions presented in this paper are those of the author(s) and do not necessarily reflect the views of the NIH or the U.S. Department of Health and Human Services.

## Author Contributions

J.Z. performed experiments, analysed data and wrote the draft of the manuscript. R.M.H. conducted computational analyses of unsupervised behavior quantification data and neuronal activity in behavioral embedding space and co-wrote the draft of the manuscript. G.C. and J.E.L. advised on the data analysis and interpretation. A.B.P. assisted with AAV construction, managed mouse colonies and edited the manuscript. J.E.L. and G.C. provided feedback and supervised the study. J.Z., G.C. and J.E.L. edited and revised the final manuscript.

---

**Extended Data Figure 1.**
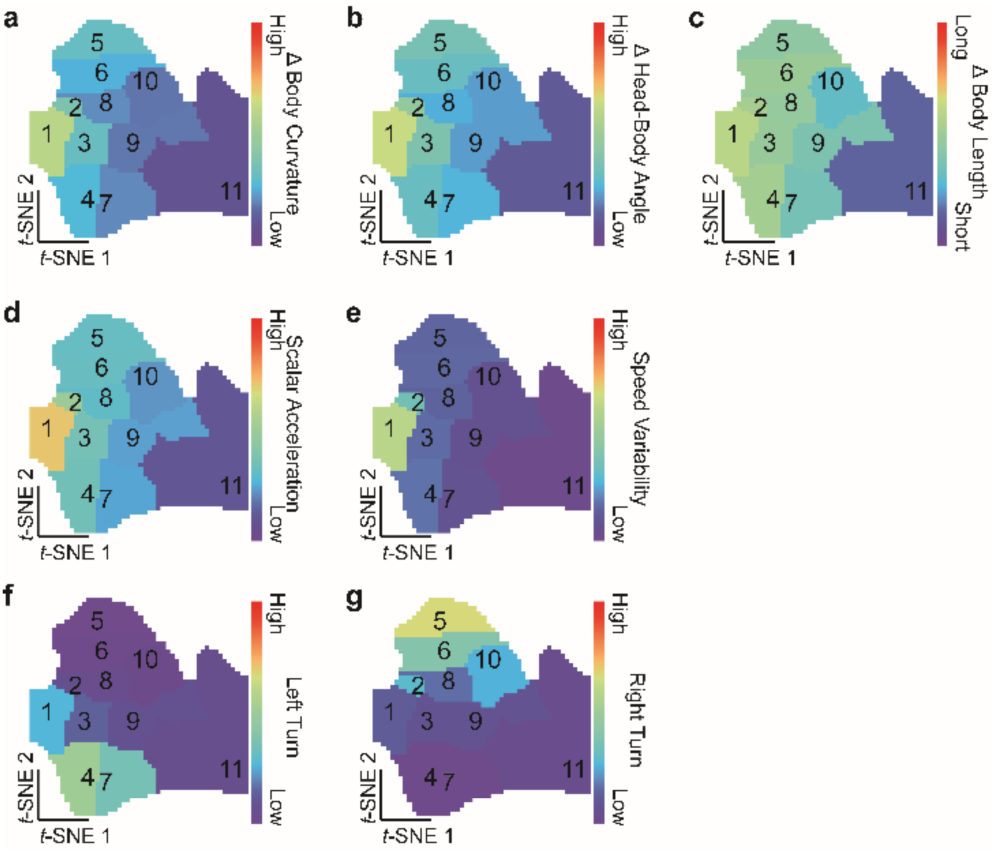
*t*-SNE clusters of locomotion features (Related to Figure 1). Average raw feature value for change in body curvature (**a**), change in head-body angle (**b**), change in body length (**c**), scalar acceleration (**d**), speed variability (**e**), windowed left-turn rate (**f**), and windowed right-turn rate (**g**).

**Extended Data Figure 2.**
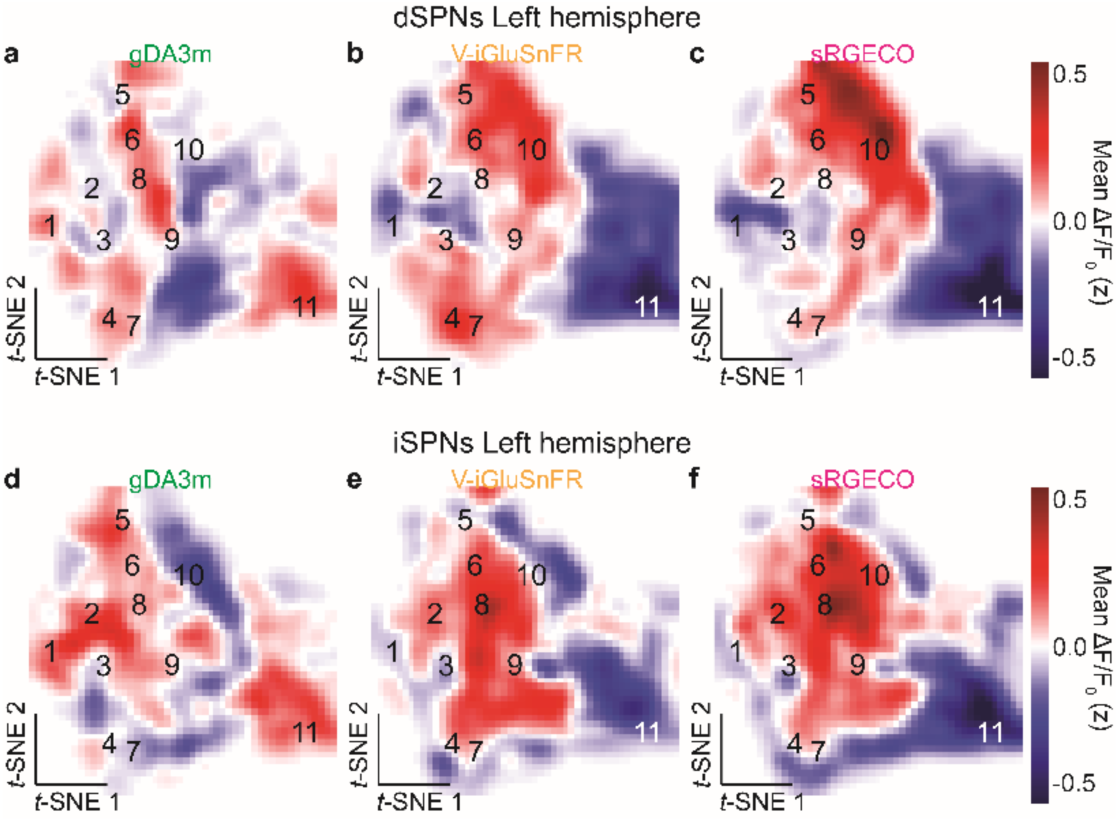
Neural signal signatures of behavior clusters (Related to Figure 1). **a-c,** gDA3m (**a**), Venus–iGluSnFR (**b**), sRGECO (**c**) across *t*-SNE behavior space in left-hemisphere dSPNs. **d-f,** gDA3m (**d**), Venus–iGluSnFR (**e**), sRGECO (**f**) across *t*-SNE behavior space in left-hemisphere iSPNs.

**Extended Data Figure 3.**
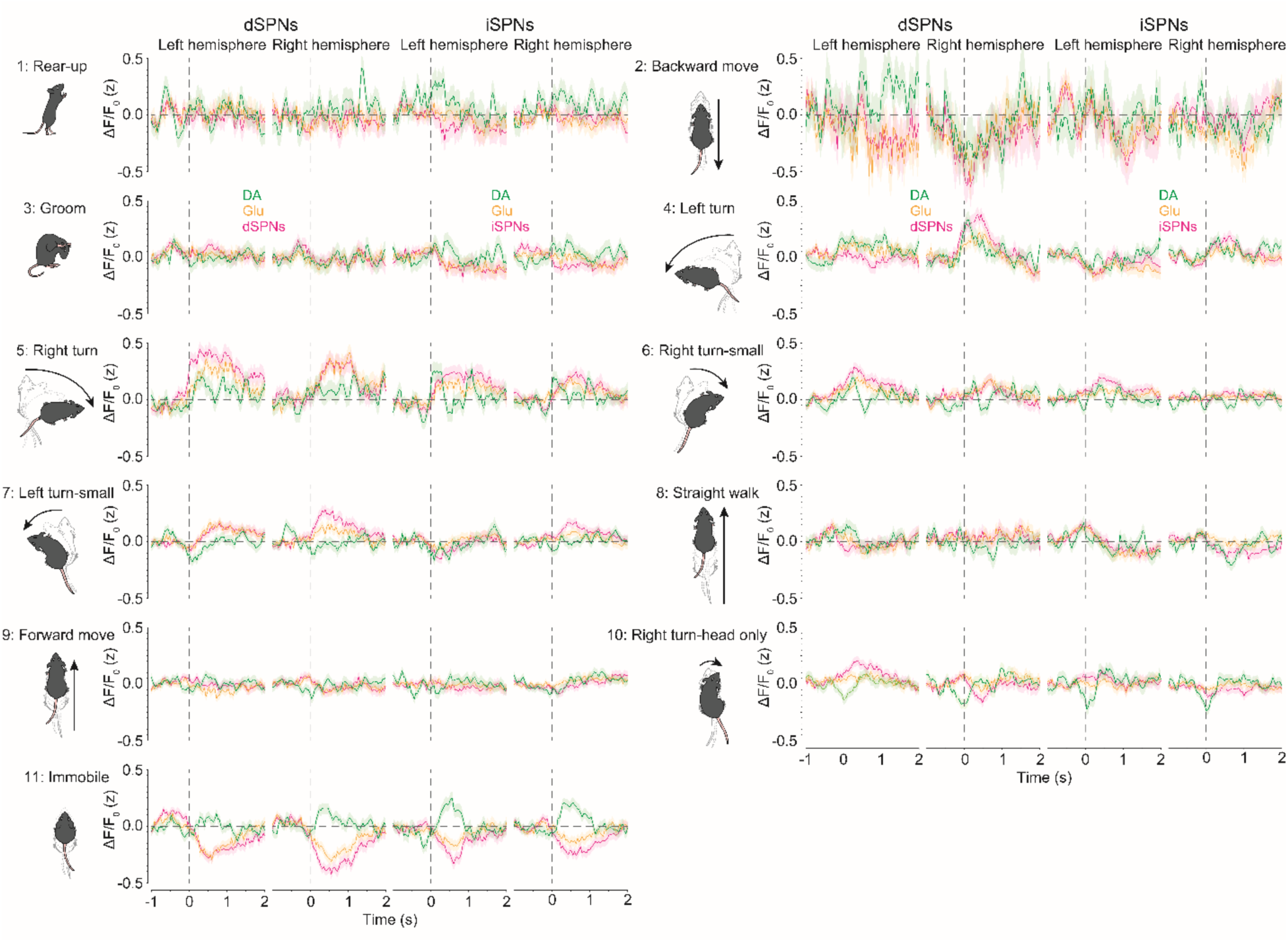
Neural signal PETHs of behavior clusters (Related to Figure 1). Averaged gDA3m, Venus–iGluSnFR and sRGECO signal from left- and right-hemisphere dSPNs and iSPNs (from left to right) aligned to onset of cluster #1 (rear-up), #2 (backward movement), #3 (groom), #4 (left turn), #5 (right turn), #6 (right-turn-small), #7 (left turn-small), #8 (straight walk), #9 (forward movement), #10 (right turn-head only), and #11 (immobile).

**Extended Data Figure 4.**
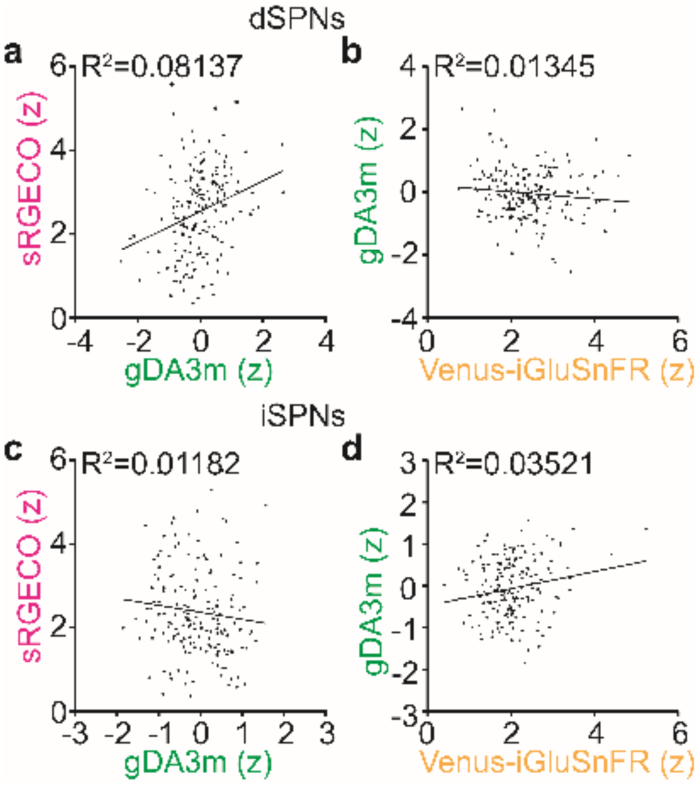
Dopamine shows no linear relationship with either neural activity or glutamate (Related to Figure 2). **a,c,** Linear regression of the magnitude of gDA3m and sRGECO in dSPNs (**a**) and iSPNs (**c**). **b,d,** Linear regression of the magnitude of Venus–iGluSnFR and gDA3m in dSPNs (**b**) and iSPNs (**d**).

**Extended Data Figure 5.**
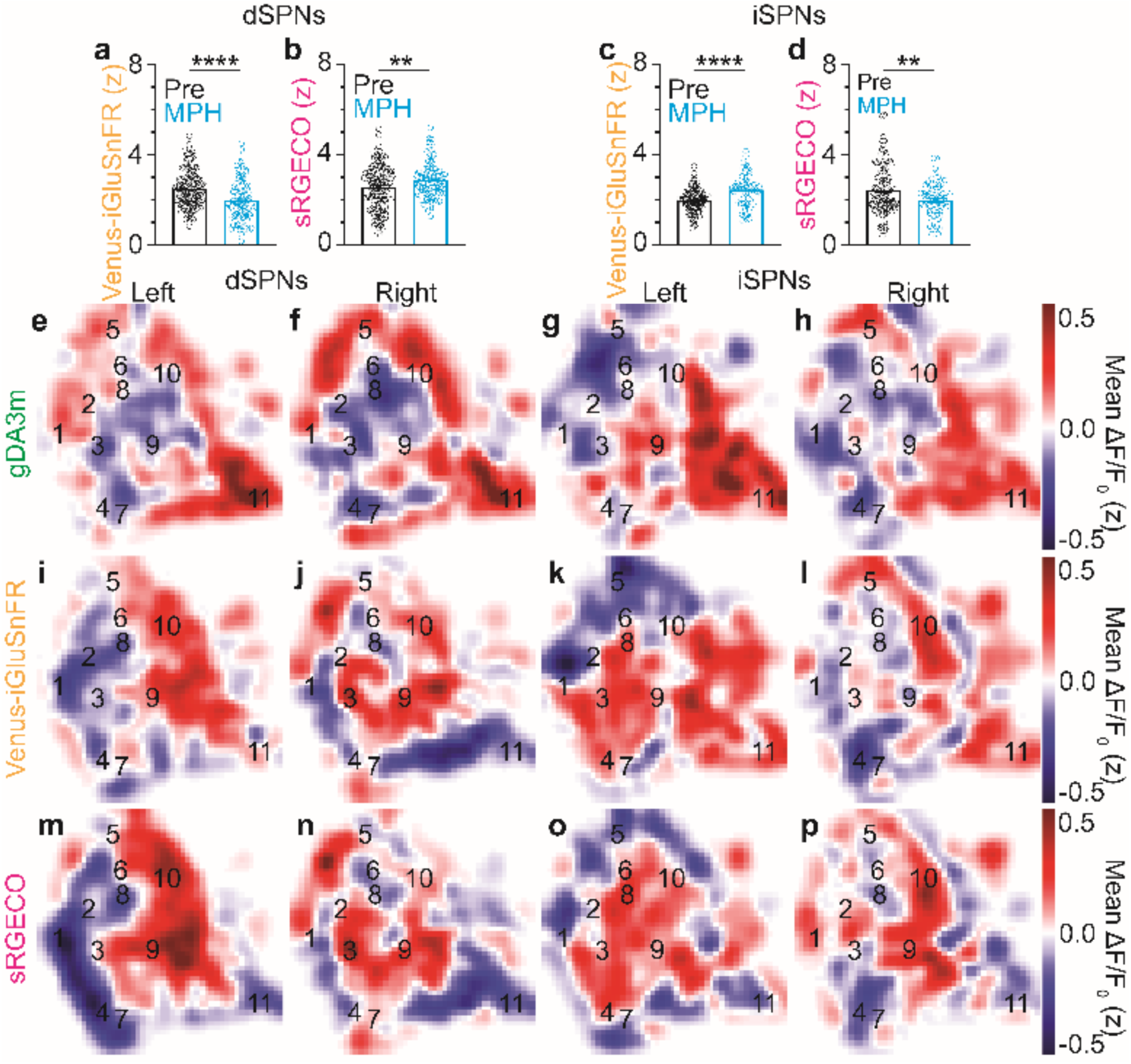
Dopamine reuptake blockade bidirectionally changes glutamate and neural activity of dSPNs and iSPNs (Related to Figure 3). **a,c,** Comparison of Venus–iGluSnFR before (black) and after MPH (blue) from dSPNs (**a**) and iSPNs (**c**). **b,d,** Comparison of sRGECO before (black) and after MPH (blue) from dSPNs (**b**) and iSPNs (**d**). **e-h,** Mean post-MPH gDA3m signal across *t*-SNE behavior space in left-hemisphere dSPNs (**e**), right-hemisphere dSPNs (**f**), left-hemisphere iSPNs (**g**), right-hemisphere iSPNs (**h**). **i-l,** Mean post-MPH Venus– iGluSnFR signal across *t*-SNE behavior space in left-hemisphere dSPNs (**i**), right-hemisphere dSPNs (**j**), left-hemisphere iSPNs (**k**), right-hemisphere iSPNs (**l**). **m-p,** Mean post-MPH sRGECO signal across *t*-SNE behavior space in left-hemisphere dSPNs (**m**), right-hemisphere dSPNs (**n**), left-hemisphere iSPNs (**o**), right-hemisphere iSPNs (**p**). Events included in the analysis: 128 for Pre-MPH and 95 for Post-MPH from dSPNs in **a,b**; 112 for Pre-MPH and 61 for Post-MPH from iSPNs in **c,d**. **, p < 0.01; ****, p < 0.0001, unpaired *t* test in **a-d**.

**Extended Data Figure 6.**
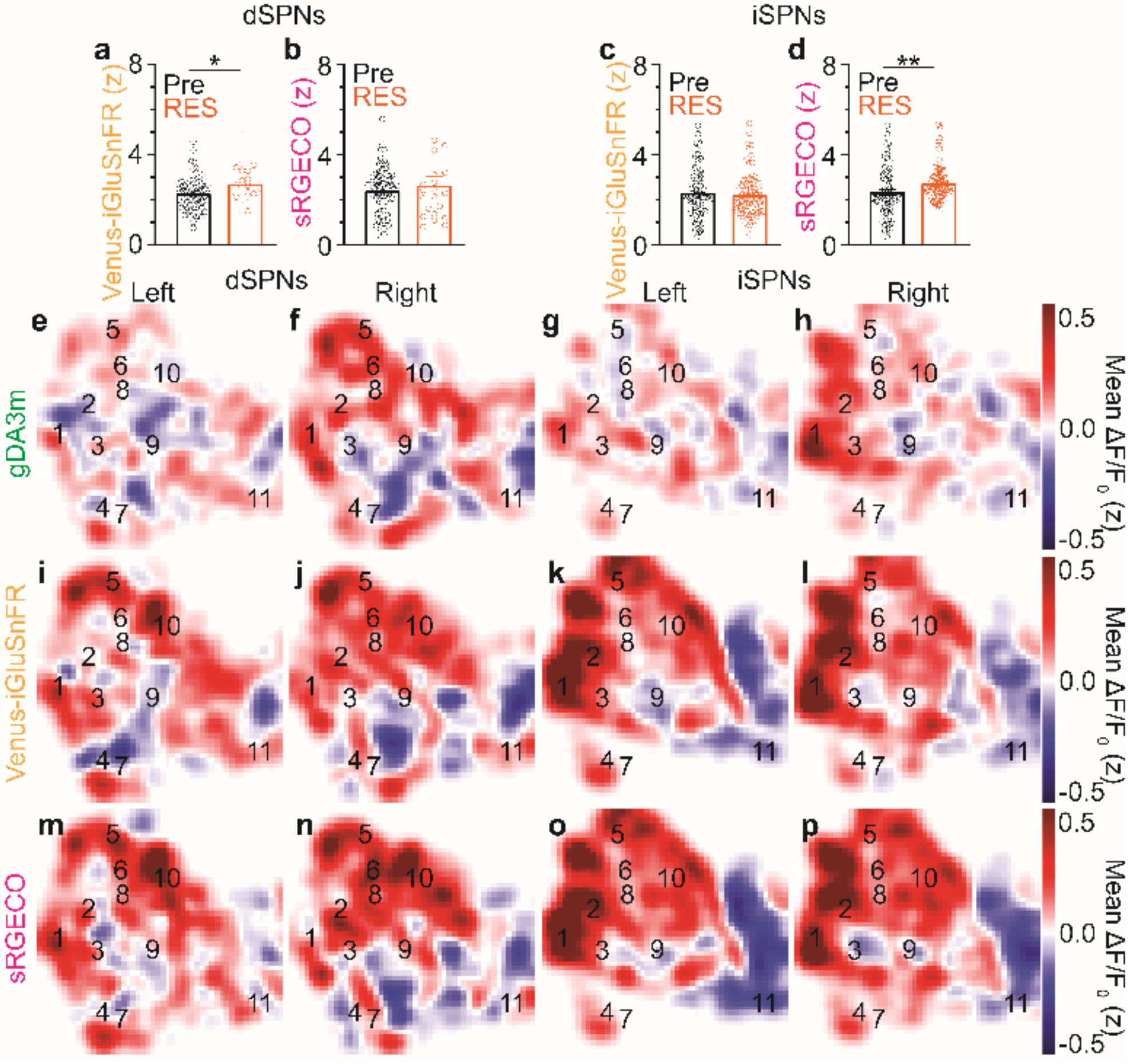
Dopamine depletion increases glutamate in dSPNs and neural activity of iSPNs (Related to Figure 4). **a,c,** Comparison of Venus–iGluSnFR before (black) and after RES (orange) from dSPNs (**a**) and iSPNs (**c**). **b,d,** Comparison of sRGECO before (black) and after RES (orange) from dSPNs (**b**) and iSPNs (**d**). **e-h,** Mean post-RES gDA3m signal across *t*-SNE behavior space in left-hemisphere dSPNs (**e**), right-hemisphere dSPNs (**f**), left-hemisphere iSPNs (**g**), right-hemisphere iSPNs (**h**). **i-l,** Mean post-RES Venus– iGluSnFR signal across *t*-SNE behavior space in left-hemisphere dSPNs (**i**), right-hemisphere dSPNs (**j**), left-hemisphere iSPNs (**k**), right-hemisphere iSPNs (**l**). **m-p,** Mean post-RES sRGECO signal across *t*-SNE behavior space in left-hemisphere dSPNs (**m**), right-hemisphere dSPNs (**n**), left-hemisphere iSPNs (**o**), right-hemisphere iSPNs (**p**). Events included in the analysis: 68 for Pre-RES and 21 for Post-RES from dSPNs in **a,b**; 67 for Pre-RES and 85 for Post-RES from iSPNs in **c,d.** n.s., p > 0.05; *, p < 0.05; **, p < 0.01, unpaired *t* test in **a-d**.

